# Differential effects of farming practice on cuckoo bumblebee communities in relation to their hosts

**DOI:** 10.1101/774406

**Authors:** Charlotte E. Howard, Alexander J. Austin, James D. J. Gilbert

## Abstract

1. Bees are important for vital pollination of wild and crop plants, but are in decline worldwide. Intensification of agriculture is a major driver of bee decline. Organic farming practices are designed to limit environmental impacts of agriculture and can increase bee abundance and species diversity. However, studies have been heavily focused towards some guilds of bees, overlooking others. This includes social brood parasites, cuckoo bumblebees, an understudied bee lineage. Little is known about bumblebee host and cuckoo population dynamics, and the effects of farming practice on cuckoo bumblebees have never previously been evaluated.
2. To compare the effects of farming practice (organic vs conventional) on the abundance, species diversity, and community dissimilarity of cuckoo bumblebees and their hosts, we compared host and cuckoo community metrics across ten matched pairs of organic and conventional farms in Yorkshire, UK.
3. As found by many previous studies, host bumblebees were more abundant on organic farms than on conventional farms. Despite this, cuckoo bumblebees were equally abundant on both farm types. Contrary to prediction, community dissimilarity and species diversity were unaffected by farm type for both host and cuckoo communities.
4. **Synthesis and applications:** Results suggest that cuckoo bumblebee community metrics are not solely driven by host community metrics, and that cuckoos may respond differently from their hosts to differences among farming practices. This could, in turn, indicate that a unified management practice is not sufficient to conserve all bumblebee species.

## INTRODUCTION

Human induced habitat degradation has led to worldwide biodiversity declines (Collen et al., 2009). For example, intensive farming practices have led to landscape simplification (Adhikari et al., 2019), increased pesticide applications (Whitehorn et al., 2012), and reductions in non-crop plant diversity and semi-natural habitat (Goulson et al., 2015). Sustainable farming systems, such as organic farming, are designed to reduce the environmental impacts of agriculture and aim to reduce biodiversity declines (Hole et al., 2005). Organic farming is less intensive (Kennedy et al., 2013; Hole et al., 2005; Power et al., 2012; Whitehorn et al., 2012), tending to use smaller sized crop fields (Kennedy et al., 2013; Hole et al., 2005), alternatives to agrochemicals, and sympathetic management of non-crop habitat, such as field margins (Hole et al., 2005; Adhikari et al., 2019). However, studies investigating the effects upon biodiversity of less intensive farming practices, such as organic farming, have often given mixed results which vary between taxa, habitat types, and landscapes (Aude et al., 2004; Bengtsson et al., 2005; Hole et al., 2005; Fuller et al., 2005; Holzschuh et al., 2008; Le Féon et al., 2010; Brittain et al., 2010). The reasons behind these differences are still poorly understood, with few studies investigating this in detail (e.g. Fuller et al., 2005; Tuck et al., 2014).

Pollinators are economically and ecologically important as they provide crucial pollination services to both wild and crop plants (Power and Stout, 2011; Scriven et al., 2015; Goulson et al., 2015; Adhikari et al., 2019), but, as with many other animal groups, are vulnerable to landscape change (Hole et al., 2005; Ceballos et al., 2015). Modern agricultural intensification plays a major role in the decline of bees, some of the most important pollinators (Holzschuh et al., 2007; Holzschuh et al., 2008; Le Féon et al., 2010; Power and Stout, 2011; Adhikari et al., 2019). This creates large expanses of resource-poor areas for bees (Carvell et al., 2006; Ollerton et al., 2014; Goulson et al., 2015; Vaudo et al., 2015; Tscharntke et al., 2012; Vasseur et al., 2013; Henckel et al., 2015), which are wholly dependent on plants for their nutritional, energetic, and developmental requirements (Goulson et al., 2011).

Organic farming is sometimes associated with more diverse, species rich, and abundant bee communities compared to conventional farming. This difference has been attributed to increased flower cover and availability of floral resources, such as non-crop plants (Aude et al., 2004; Holzschuh et al., 2007; Rundlöf et al., 2008; Le Féon et al., 2010; Power and Stout, 2011). These are essential for bee nutrition (Brodschneider and Crailsheim, 2010; Whitehorn et al., 2012; Di Pasquale et al., 2013; Adhikari et al., 2019). However, studies have often grouped all bee guilds together (Holzschuh et al., 2007; Holzschuh et al., 2010; Power and Stout, 2011) or studied guilds in isolation (Wintermantel et al., 2019; Nooten and Rehan, 2019). The extent to which organic farming practices can benefit bees varies between guilds; for example among honeybees, social bumblebees, and solitary bees (Le Féon et al., 2010; Holzschuh et al., 2008), and at species-level within the *Bombus* genus (Rundlöf et al., 2008). It has been suggested that this is due to varied foraging distances (Holzschuh et al., 2008; Rundlöf et al., 2008) and nesting resource requirements of different bee guilds (Rundlöf et al., 2008). Given bees’ highly varied ecology (Goulson, 2003; Goulson and Darvill, 2004; Lhomme and Hines, 2018), the assumption that all bee guilds will be affected in the same way by farming and management practices may therefore not be valid (Rundlöf et al., 2008). To our knowledge there have been very few studies of the mechanism underlying differential effects of farming practices upon different guilds (e.g. Kennedy et al., 2013), in line with a paucity of such studies generally.

To date, studies investigating the effects of organic farming practices have been heavily focused on specific bee groups, such as honeybees and social bumblebees (Serrano and Guerra-Sanz, 2006; Velthuis and Van Doorn, 2006; Holzschuh et al., 2008; Le Féon et al., 2010; De Luca et al., 2013). As a result, some bee guilds in particular are underrepresented in the literature (Wood et al., 2016; Lhomme and Hines, 2018). Among these underrepresented guilds, obligate social brood parasites, cuckoo bumblebees, are amongst the least well studied (Power and Stout, 2011; Waters et al., 2011; Lhomme and Hines, 2018). Cuckoo bumblebees are a monophyletic subgenus of the *Bombus* genus, called *Psithyrus*, yet despite being closely related, their ecology varies a great deal from that of their host (Cameron et al., 2007; Lhomme and Hines, 2018). Unlike other bumblebees, *Psithyrus* species have lost the ability to produce workers and collect pollen (Lhomme and Hines, 2018). Emergence of cuckoo bumblebees from dormancy is delayed in order to allow time for the host to establish a colony and produce a worker generation (Lhomme and Hines, 2018). A cuckoo queen sneaks into a host colony, matching her chemical signals to those of the target nest to avoid detection by the host workers (Lhomme and Hines, 2018). Following this, the cuckoo attacks the host queen, most often killing her, then ejects the host eggs from the nest and enslaves the host workers to care for the parasitic brood (Lhomme and Hines, 2018). *Psithyrus* species are important pollinators, and evidence suggests that they are more at risk of extinction than their hosts (Kosior et al., 2007; Suhonen et al., 2015; Lhomme and Hines, 2018).

Host-parasite relationships in hymenopterans, such as bees, can be influenced by factors such as nutrition (Goodell, 2003; Di Pasquale et al., 2013; Maure et al., 2016), pesticides (Pettis et al., 2012), and host availability (Mahmoudi et al., 2010; Kaspi et al., 2011), which can affect parasitism success rates, and host recovery (Goodell, 2003; Di Pasquale et al., 2013; Maure et al., 2016) among other effects. Since all these factors can be affected by farming practice, it is likely that farming practice can mediate the host-parasite relationship between cuckoo and host bumblebees. Despite this, the effect of farming practice on cuckoo bumblebee diversity and abundance has never previously been investigated.

In this study we compare the effects of farming practice, organic and conventional, on the community metrics of cuckoo bumblebees and their hosts, using captured samples of bumblebees from ten matched pairs of organic and conventional farms in Yorkshire. Since farming practice can affect the abundance, diversity, and species richness of bumblebees (Rundlöf et al., 2008; Adhikari et al., 2019), it can be expected that this could also cause knock-on effects to the cuckoo bumblebee community (Arneberg et al., 1998; Steffan-Dewenter and Schiele, 2008; Suhonen et al., 2015). Firstly, we predicted that species richness, diversity, and abundance of hosts, and cuckoos, will be higher on organic farms than on conventional farms. Obligate brood parasites such as cuckoo bumblebees, are entirely dependent on their hosts for reproduction, thus the host can be considered a resource (Steffan-Dewenter and Schiele, 2008; Suhonen et al., 2016). Therefore, elements of their community structure are likely to be driven by those of their host (Murray et al., 2009; Suhonen et al., 2015; Suhonen et al., 2016). Therefore, we also predicted that cuckoo bumblebee community metrics would be driven by the community metrics of their hosts.

## METHODS

### Study sites

Ten organic farms were identified in the East Riding of Yorkshire and North Yorkshire, UK. Sampling had a paired design to reduce confounding factors due to variation in the local environment (Holzschuh et al., 2008; Holzschuh et al., 2010; Kennedy et al., 2013). Organic and conventional farms were paired together based on crop type, soil type, and locality (Power et al. 2012) (see Figure 1 for farm locations), since these factors can influence the effects of farming practice (Rundlöf et al., 2008; Power and Stout, 2011; Tuck et al., 2014). All paired sites were at least four kilometres apart in order to minimise the risk of any bees foraging at both locations (Zurbuchen et al. 2010). Sample collection took place in the arable portion of each farm, except on the two pairs of pastoral farms where sampling took place within the pasture. A single field was chosen as the sampling site for each area. Crop or crop type of the sampling area was matched for each pair.

**Figure 1:**
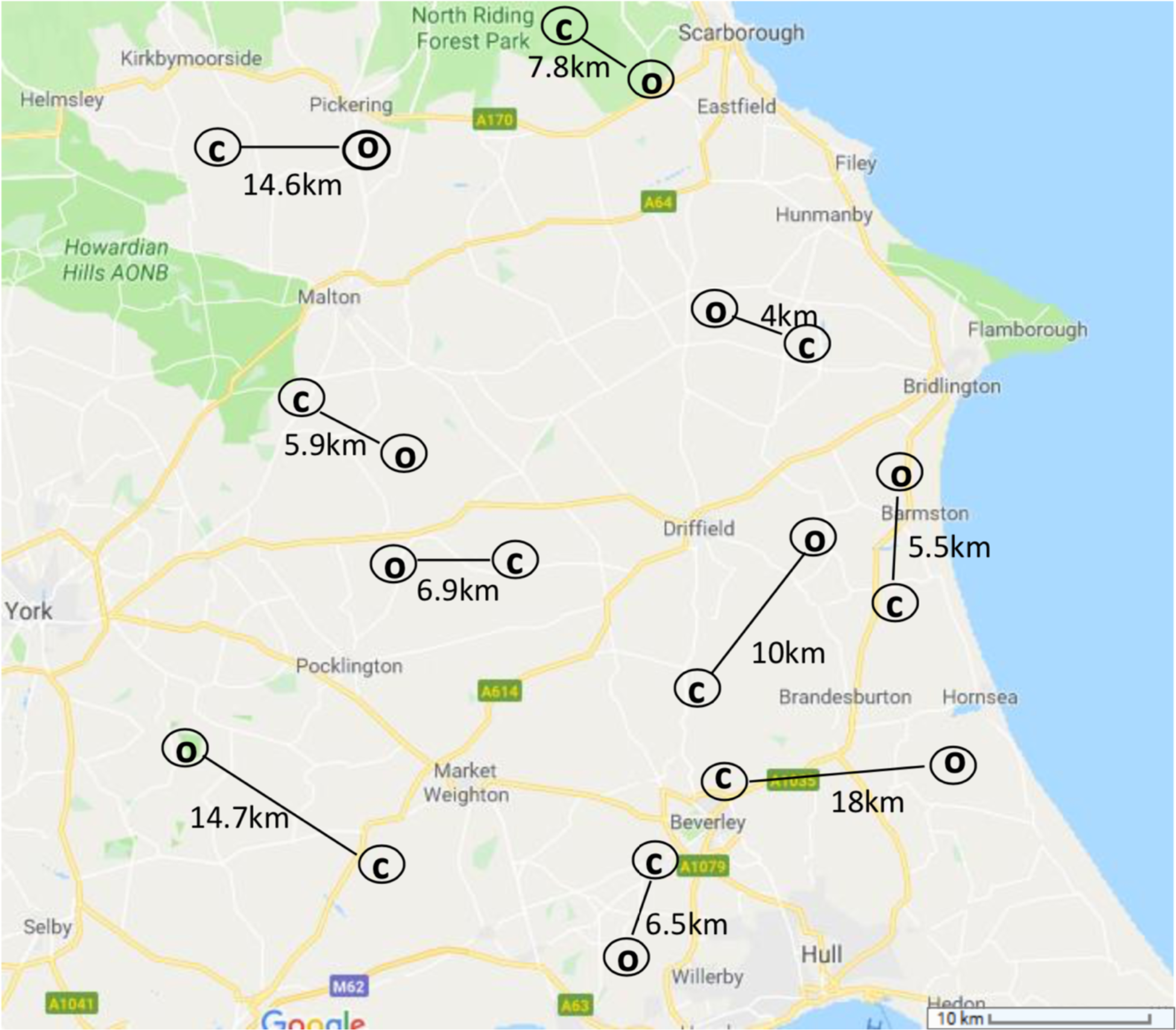
Study sites for transect sweep-net sampling of bees in East Riding of Yorkshire and North Yorkshire, UK. Each site location is marked with a circle. Eight of the ten farm pairs were arable or mixed arable/pastoral farms, two pairs were dairy/beef farms. Cereals were the primary crop at all farm sites. Farms were matched by locality and soil type and were at least 4km apart to minimise risks of any individuals foraging at both sites.

### Sample collection

Samples were collected from May to September, 2016, between 09:00 and 16:00, on dry, sunny days with wind speeds up to moderate (Forup and Memmott, 2005; Power and Stout, 2011). There were single hour-long visits to each site per month. This spread the sample collection, minimising the effects on the environment and bee communities. Each sampling site was visited five times in total. The order in which each farm within a pair was visited was alternated every sampling session to ensure that a) farms within pairs were sampled at the same time of day, and b) each farm was sampled during both the morning and afternoon (Power and Stout, 2011). These methods reduced the effects of confounds such as weather and time of day. At each visit of every site, sampling was carried out using a sweep net along a transect line, pausing the stopwatch for sample processing (ensuring a full hour of sampling effort). The transect was ‘E’ shaped, with the spine (20×1m) running along the hedgerow/margin and the three arms (20×1m) stretching into the crop. This was walked at a constant pace. This shape was chosen in order to sample a representative amount of each habitat type; hedgerow, margin and crop. For example, as the largest proportion of the fields were the crop, the largest proportion of sampling time was spent here. The location of the first transect at each site was decided with a random number generator, and subsequent visits were done in a clockwise fashion, with each margin sampled at least once. All bees were stored individually in a 50ml falcon tube in a cool bag with ice to reduce stress. Bees were euthanised by placing into a freezer at −20°C, and stored at −80°C. Bees were thawed in the fridge before manual identification to species level under a light microscope using Falk (2015).

Manual identification was used since the sample size was too large to use molecular techniques for the entire sample. However, cuckoo bumblebee species and some host bumblebee species are difficult to distinguish morphologically (lucorum complex: Bossert et al., 2016; Psithyrus spp.: Falk, 2015). Therefore, DNA barcoding (Supplementary Methods) of a subset of samples was used to confirm the accuracy of the manual identification to improve confidence in the accuracy of the data used for the community statistics.

### Community statistics

Community statistics were carried out using R software (R Core Team, 2017) with the following packages; iNEXT (Hsieh et al. 2019), lme4 (Bates et al. 2015). This included calculation of species richness, abundance, and Simpson’s diversity index. Simpson’s diversity index was chosen as it is a good general-purpose statistic (Magurran, 2003) and is a more compatible index to use with the iNEXT package than the Shannon’s index. Sample-size integrations of rarefaction and extrapolation of data with ‘iNEXT’ (Appendix 2) were used to standardise and compare species richness and diversity of hosts and cuckoos across farm type (Hsieh et al., 2016). Generalised linear mixed models (GLMMs) were used to assess the effects of farm type and month on cuckoo (n=89) and host (n=1,247) community metrics. Separate models were fitted with ‘abundance’, ‘species richness’ and ‘Simpson’s diversity index’ as response variables, respectively. Each model had ‘farm type’ and ‘visit month’ as predictors, and ‘farm pairs’ as a random effect. A non-metric multidimensional scaling ordination (NMDS) plot on Bray-Curtis distance metrics was used to visualise community dissimilarity.

## RESULTS

Out of the sample tested, DNA barcoding confirmed accurate identification of all host individuals except for two (96%, n=50) and all cuckoo bumblebees but 5 (94%, n=87). One sample matched to a species of water mould, *Peronospora flava*, and was discarded due to contamination.

### Abundance

We found 8 species of host bumblebee (n=1,247), along with 7 species of cuckoo bumblebee (n=89). Abundance of hosts and cuckoos varied over the months in the season (Figure 2B). For example, on organic farms, abundance of hosts peaked in July, but on conventional farms, appeared to increase linearly from May to September (GLMM containing farm type and visit month as predictors with farm pairs as a random factor, dropping the farm:month interaction, *χ*^2^_4,16_=10.83, p< 0.05; dropping bee type:month, *χ*^2^_4,16_=35.12,p<0.001; Figure 2B). Total abundance of cuckoos did not vary across farm type. Abundance of hosts was higher on organic farms than conventional farms (Figure 2A). Cuckoo abundance was driven mainly by two months in particular, July and August. Abundance of cuckoos was higher on organic farms in July, and higher on conventional farms in August, despite host abundance being lower on conventional farms in August. Cuckoos were less abundant than hosts in all five months (Figure 2B).

**Figure 2:**
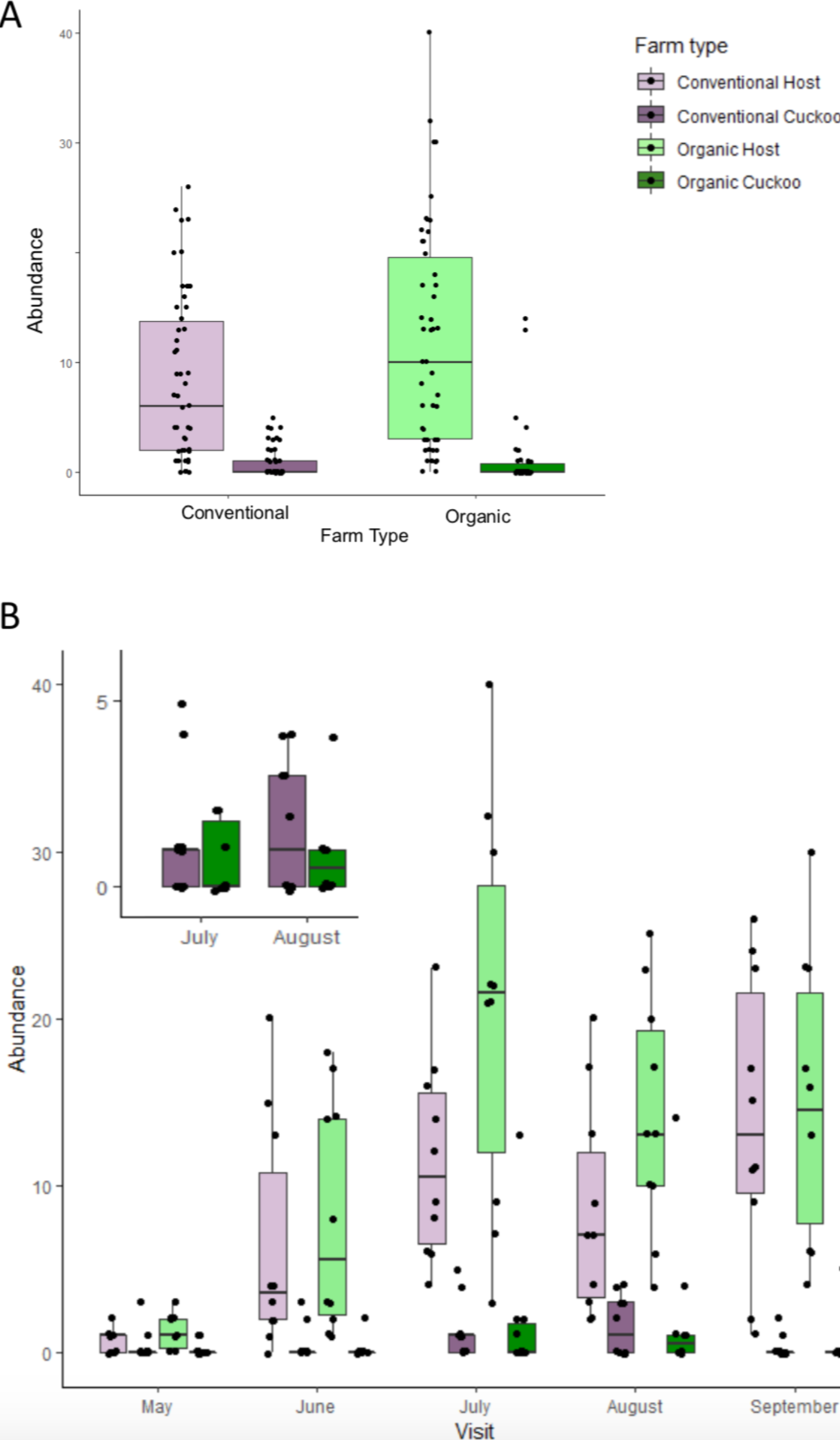
Abundance of cuckoo and host bumblebees found on organic and conventional farms in total (A) and by month (B). For clarity, a larger-scaled section of cuckoo bumblebee abundance across the months is shown in the top left of graph B for their two key months of activity, July and August.

### Species richness and Simpson’s diversity index

Species richness and Simpson’s diversity index broadly reflected similar patterns through the months of the season for hosts and cuckoos (Figure 3A and 3C). Total richness and diversity of hosts communities were similar on organic and conventional farms, and the same was true for cuckoo communities (Figure 3B and 3D). Species richness and Simpson’s diversity index differed between hosts and cuckoos, and changed across the months (species richness: GLMM containing farm type and visit month as predictors with farm pairs as a random factor, dropping the main effect of visit month, χ^2^_4,8_=25.115, P<0.001; and bee type, χ^2^_1,9_=24.033, P<0.001) (Simpson’s diversity index: GLMM containing farm type and visit month as predictors with farm pairs as a random factor, dropping the main effect of visit month, χ^2^_4,8_=24.087, P<0.001; and bee type, χ^2^_1,8_=10.335, P=0.001; Figure 3). Temporal patterns in richness and diversity did not differ between farm types. On conventional farms, cuckoo richness and diversity peaked in August (Figure 3A and C). In contrast, hosts were most species-rich in July, but were more diverse in August (Figure 3A and 3C). On organic farms, richness and diversity of both hosts and cuckoos peaked in July (Figure 3A and 3C). In total across the whole season, hosts were more diverse and species-rich than the cuckoos (Figure 3B and 3D).

**Figure 3:**
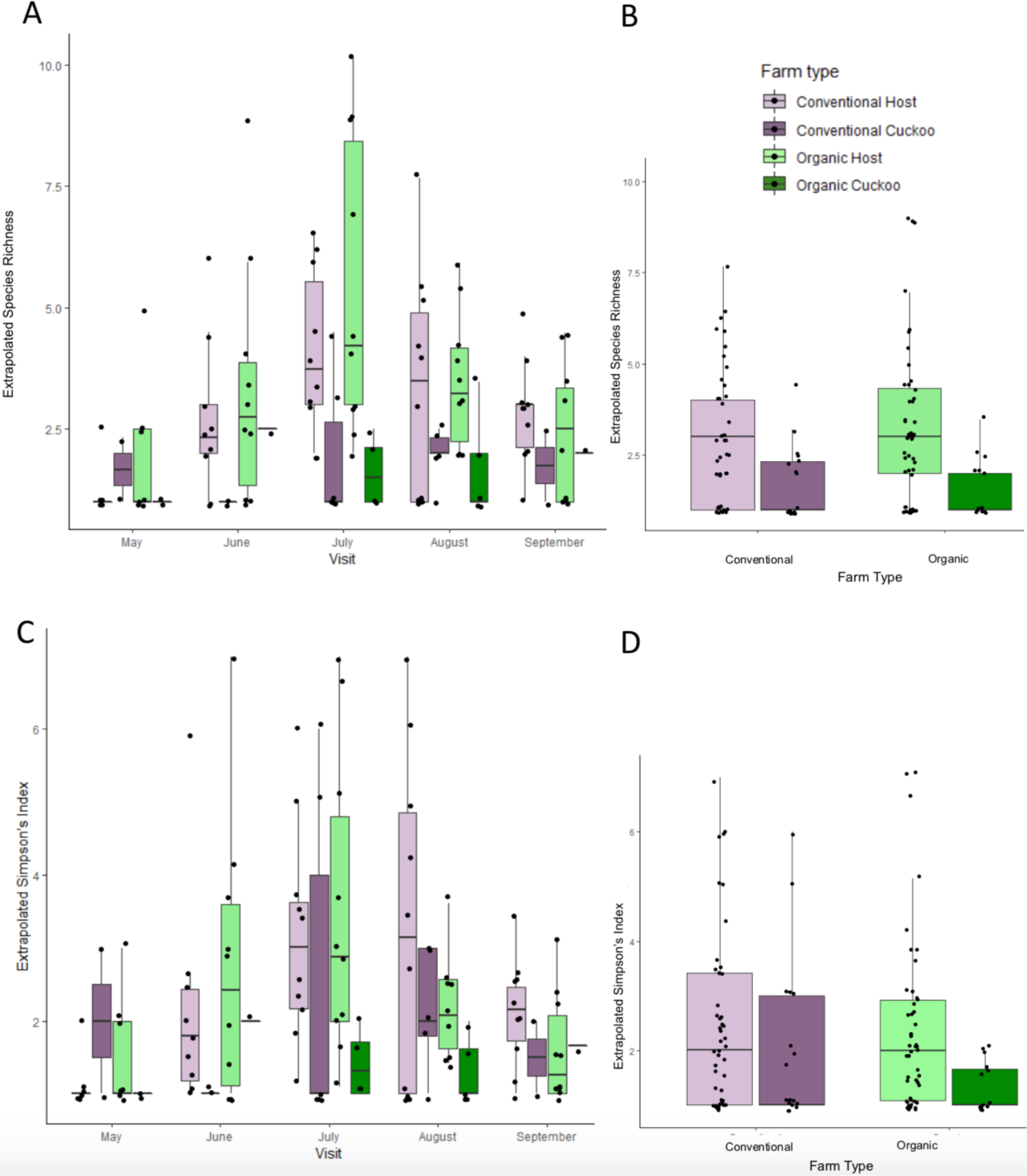
Extrapolated species richness of cuckoo and host bumblebees found on organic and conventional farms by month (A), and in total (B). Extrapolated Simpson’s diversity index of cuckoo and host bumblebees found on organic and conventional farms by month (C), and in total (D). Sample-size integrations of rarefaction and extrapolation of data with ‘iNEXT’ (see text) were used to standardise and compare species richness and diversity.

### Community Dissimilarity

The NMDS plot shows that the species composition was very similar on organic farms and conventional farms for all bumblebees, both hosts and cuckoos (Figure 4). For cuckoos, there was almost a complete overlap of the bumblebee communities at each farm type, as with the host communities (Figure 4). Cuckoo communities on organic farms showed a very small amount of variation compared to cuckoo communities on conventional farms (Figure 4B). The species composition of cuckoos on conventional farms was a subset of the species composition of cuckoos on organic farms (Figure 4B). Splitting the data by month revealed no particular patterns (data not shown).

**Figure 4:**
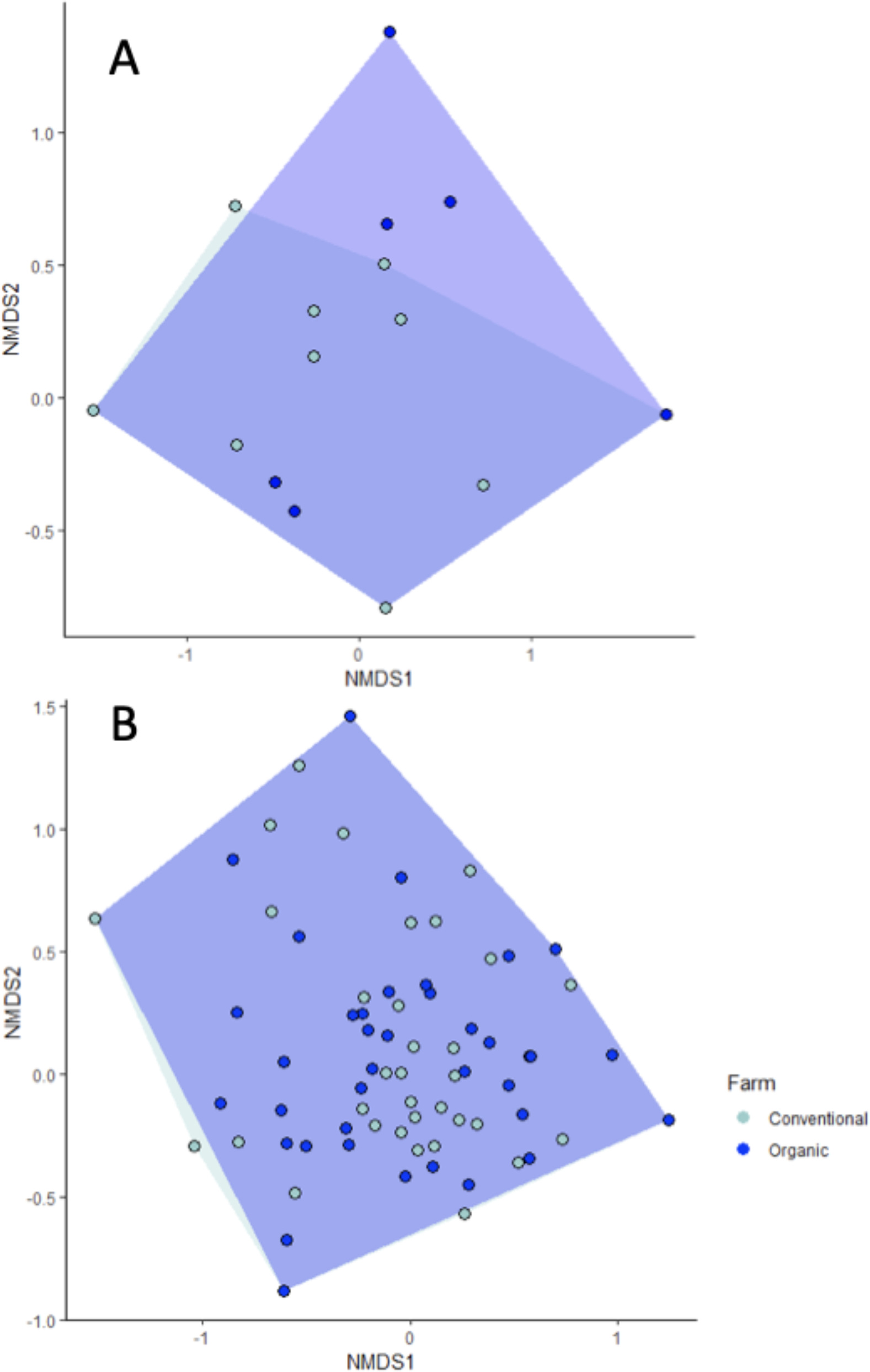
A non-metric multidimensional scaling ordination (NMDS) plot on Bray-Curtis distance metrics to visualise community dissimilarity of A) cuckoo bumblebees, B) host bumblebees, collected from organic and conventional farms. Dark blue area shows communities on organic farms, and the pale blue area shows communities on conventional farms.

## DISCUSSION

There have been few previous studies that investigate why organic farming should not be universally beneficial to farmland wildlife; and none investigating the effect of farm type on the community structure of cuckoo bumblebees (Power and Stout, 2011; Lichtenberg et al., 2017). Interestingly, and contrary to prediction, cuckoo abundance was *not* primarily driven by the abundance of their hosts. As predicted, hosts were more abundant on organic farms than on conventional farms,but despite this, cuckoos were equally abundant across both farming practices. In both hosts and cuckoos, both diversity and species richness were similar on organic and conventional farms; both metrics showed broadly similar patterns across the months within the season, and the two guilds showed very similar species composition across farm types.

Perhaps the reason why cuckoos were equally abundant on both organic and conventional farms, despite their hosts being more abundant on organic farms, was due to decreased health and nutrition of hosts on conventional farms, and hence increased susceptibility to cuckoo usurpation. For example, the increased use of pesticides on conventional farms could have negative effects on bee health; pesticides, for instance, can impair immune response (Pettis et al., 2012), colony growth (Rundlöf et al., 2015), queen mortality (Scholer and Krischik, 2014), susceptibility to parasites (Pettis et al., 2012), and foraging behaviour (Yang et al., 2008; Henry et al., 2012). Hosts emerge earlier from overwintering than cuckoos, hence are active for a longer period of time within the season, thus are more likely to be exposed to insecticides than cuckoos (Botías et al., 2015; Lhomme and Hines, 2018). Similarly, unlike cuckoos, the hosts will forage for pollen in addition to eating nectar (Lhomme and Hines, 2018).

Additionally, since organic farms generally offer increased diversity and abundance of floral resources (Rundlöf et al., 2008; Le Féon et al., 2010; Power and Stout, 2011), bees on organic farms can benefit from a more nutritionally diverse diet (Carvell et al., 2006; Ollerton et al., 2014; Goulson et al., 2015; Vaudo et al., 2015). Not all plants produce pollen of sufficient nutritional quality to meet the requirements of bees (Filipiak, 2017a,b). Since bee larvae require specific nutrients for growth and development (Austin & Gilbert 2018), large quantities of food can not compensate for poor quality (Filipiak, 2018). Bumblebees are wholly dependent on flowers for their nutritional, energetic, and developmental requirements (Goulson et al., 2011). In bees, nutrition affects body size, immune defence (Alaux et al., 2010; DeGrandi-Hoffman et al., 2010), fecundity, longevity, individual health, colony health, and colony productivity (Brodschneider and Crailsheim, 2010; Di Pasquale et al., 2013).

Therefore, perhaps nutrition and health are linked with a host queen’s ability to defend herself from attacks by the large, heavily armoured *Psithyrus* queen (Lhomme and Hines, 2018; Fisher and Sampson, 1992). One study has investigated this idea in cuckoo bumblebees (Cartar and Dill, 1991), and found that food scarcity increased susceptibility of bumblebee hosts to brood parasitism by *Psithyrus insularis* (Cartar and Dill, 1991). While this idea in cuckoo bumblebees has received almost no attention in the literature since, similar results have been found for other parasitic Hymenoptera (Goodell, 2003; Di Pasquale et al., 2013; Maure et al., 2016). Future studies could further investigate the effects of health and nutrition on invasion success of *Psithyrus* species, for example using controlled diets under laboratory conditions (Goodell, 2003). Such research would be critical to understanding the complex, differential ecology and communities of hosts and cuckoos.

Studies investigating the effects of organic or other less intensive farming practices on bee richness and diversity have found mixed results (Bengtsson et al., 2005; Hole et al., 2005; Le Féon et al., 2010; Aude et al., 2004; Brittain et al., 2010). The nesting strategies and specialisation of guilds means that the availability of the correct quantity and quality of resources, spatially and temporally, are the key determinants for which species a landscape can support (Tscharntke et al., 2005). Many studies have found increased species richness, and diversity of bees and bumblebees on organic farms, and as with abundance, have mainly attributed this to increased floral resources (Holzschuh et al., 2007; Holzschuh et al., 2008; Le Féon et al., 2010). However, the results of this study found no difference in species richness, diversity, or community dissimilarity across farm type. An explanation for this finding could lay in which specific host species are found in the area. The majority of host bumblebees found in the UK are generalist feeders (Falk, 2015). Hence, perhaps increased floral richness would not attract additional host species to the area, and therefore would not attract additional cuckoo species either. All of the host species found on the conventional farms were also found on the organic farms. Additionally, landscape effects must be considered (Holzschuh et al., 2008; Rundlöf et al., 2008; Weibull et al., 2000), for example, organic farming may not always show the same benefits to pollinators, if isolated within a mosaic of conventional or other farms (Brittain et al., 2010). Organic farms in Yorkshire are few, and are thus relatively isolated. Hence, perhaps the effects of organic farming practices are masked by the surrounding landscape. Future studies could include landscape effects in the analyses, for example by separating farm pairs into groups depending on habitat type (Power and Stout, 2011).

Host bumblebees were more abundant on organic farms than on conventional farms. This is supported by a wide body of literature (Aude et al., 2004; Morandin and Winston, 2005; Rundlöf et al., 2008; Le Féon et al., 2010; Power and Stout, 2011). This result is commonly assigned to reduced abundance and diversity of floral resources on conventional farms (Aude et al., 2004; Holzschuh et al., 2007; Rundlöf et al., 2008; Le Féon et al., 2010; Power and Stout, 2011; Whitehorn et al., 2012; Adhikari et al., 2019), often resulting from less intensive agricultural practices (Bengtsson et al. 2005, Hole et al. 2005, Roschewitz et al. 2005; Gabriel et al. 2006; Holzschuh et al. 2007; Holzschuh et al., 2008; Austin and Gilbert, 2019). Hosts were also more abundant than cuckoos in total across the season. Generally, macro-parasites have a lower population size than their host (Murray et al., 2009) and, in the case of cuckoo bumblebees, emerge later in the season once the hosts have already established a colony and produced workers (Erler and Lattorff, 2010; Suhonen et al., 2015; Lhomme and Hines, 2018). Similarly, cuckoo bumblebees do not produce workers, hence are naturally rarer than their hosts (Lhomme and Hines, 2018).

Cuckoo abundance across the season did not follow the same patterns as host abundance. Despite a lower abundance of hosts on conventional farms, cuckoo bumblebees were equally as abundant on organic farms as they were on conventional farms. This was not expected, since for obligate parasites such as cuckoo bumblebees, the host can be considered a resource (Suhonen et al., 2016), hence elements of their community structure are likely driven by those of their host (Murray et al., 2009; Suhonen et al., 2015; Suhonen et al., 2016). Kleptoparasites benefit from the resources gathered and constructed by their host (Murray et al., 2009), therefore the cuckoo presumably gains a fitness benefit from an increase in the resources available to their hosts. There have been very few studies on cuckoo bumblebees (Lhomme and Hines, 2018), hence future studies could shed more light into the relationship between cuckoo and host communities.

Despite receiving little attention in the literature (Williams and Osborne, 2009; Lhomme and Hines, 2018), cuckoo bumblebees are important pollinators (Kosior et al., 2007; Lecocq et al., 2011). As such, further research is needed to fully understand their ecology (Lhomme and Hines, 2018) and evidence suggests that they are even more at risk of extinction than their hosts (Suhonen et al., 2015). Whilst the ecology and community metrics of hosts will affect cuckoo community metrics due to their reproductive dependency on their host (Suhonen et al., 2016), the results suggest that perhaps cuckoos can respond differently from their hosts to environmental factors, such as farming practice. Differential effects of farming practices have previously been found among groups within the bee superfamily, for example among honeybees, bumblebees and solitary bees (Le Féon et al., 2010; Holzschuh et al., 2008) and are even found within the *Bombus* genus (Rundlöf et al., 2008). The results of this study support the idea that perhaps hosts and cuckoo bumblebees need to be considered and targeted separately when designing conservation measures and management practices. There are more than two hundred species of bumblebee (Williams et al., 2008), with 25 resident in the UK, within which there is great variation in colony size (Rundlöf et al., 2008), feeding ecology, habitat, and reproduction (Goulson, 2003; Goulson and Darvill, 2004; Lhomme and Hines, 2018). As such, the assumption that all guilds of bees will be affected in the same way by farming and management practices may not be valid, and the assumption that a unified management practice is sufficient to conserve all bumblebee species may be naïve (Rundlöf et al., 2008).

## ACKNOWLEDGEMENTS

The authors thank L. Handley for useful discussions on the manuscript and G. Sellers for help with lab work.

## Authors’ contributions

All authors initially conceived the idea for this work. JG and AA designed the overall infrastructure underpinning the study, while CH and AA designed the specifics of this investigation. CH performed all molecular work, conducted all data analysis and wrote the manuscript. AA performed all fieldwork and collected all field samples. All authors edited, contributed to and approved the final manuscript.

## Ethics Statement

This study was ethically approved by the University of Hull Faculty of Science & Engineering Ethics Committee under licence FEC_55_2017_U_FEC.

