## supplementary material for "Differential effects of farming practice on cuckoo bumblebee communities in relation to their hosts"

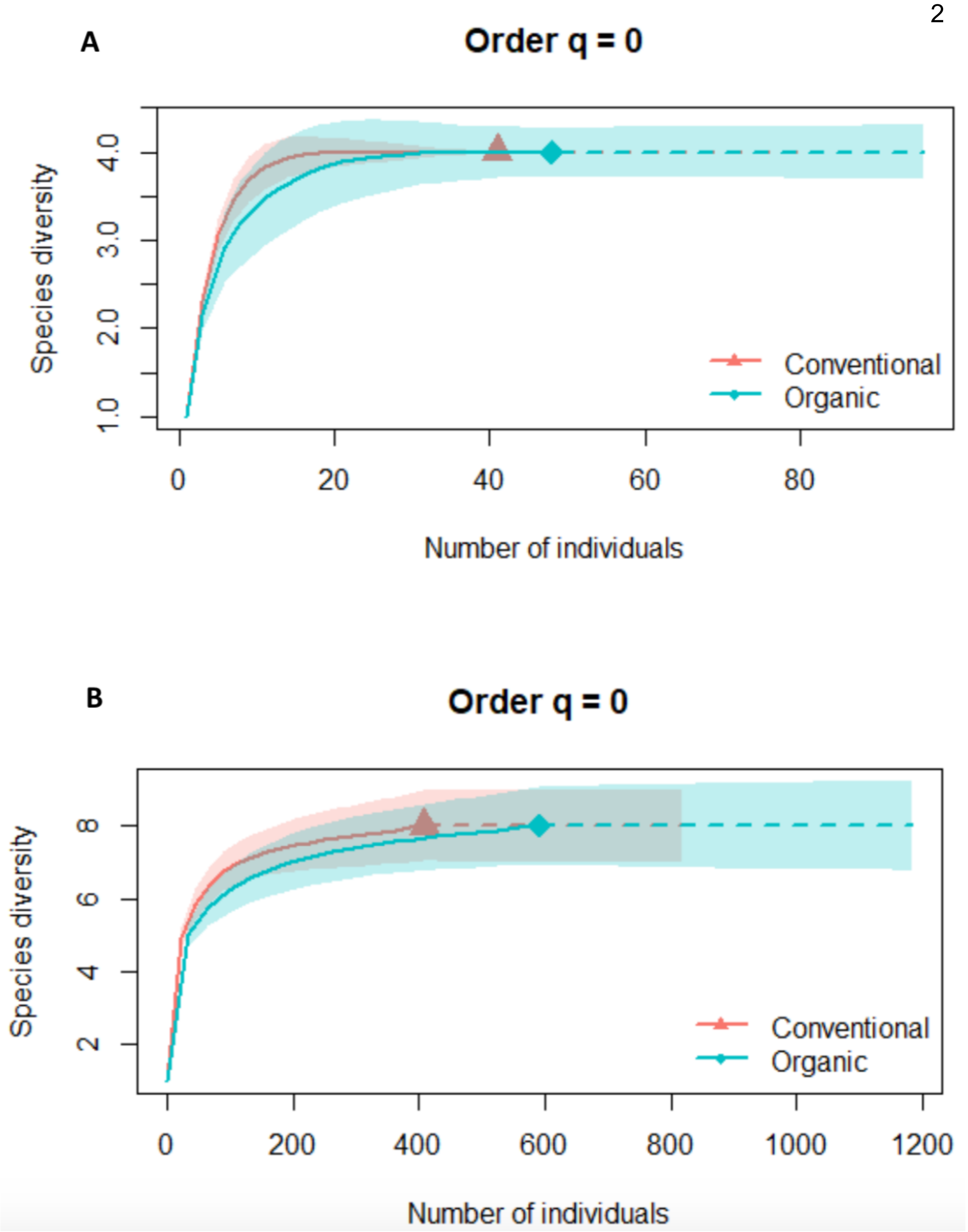


**Figure S1:** Extrapolation curve made using sample-size integrations of rarefaction and extrapolation of data with statistical R packages, iNEXT (Chao et al. 2019), lme4 (Bates et al. 2015), and ggplot2 (Wickham 2016). The curves show the extrapolation of species diversity data for cuckoo bumblebees (A), and their hosts (B). The extrapolated data were used to standardise and compare species richness and diversity.

**Supplementary Methods: DNA barcoding sequencing**

Barcoding was carried out using all cuckoo bumblebee samples (n=89) and a representative subsample their hosts (n=60). Molecular identification was carried out using the DNA barcodes (cytochrome *c* oxidase subunit 1, *COI,* Folmar fragment (658bp)) (Folmer et al., 1994) from captured bee specimens, as is standard (Williams et al., 2011; Carolan et al., 2012; Williams et al., 2012; Williams et al., 2013). DNA was extracted following the HotSHOT method (Truett et al., 2000) from clippings of the leg, antenna and tongue. PCRs were performed on an Applied Bio-systems Vertiri Thermal Cycler with the following profile: 94°C for 3 minutes, followed by 37 cycles of denaturation at 94°C for 30 seconds, annealing at 52°C for 1 minute and extension at 72°C for 1 minute and 30 seconds, with a final extension time of 10 min at 72°C. A 750bp fragment was targeted. 25µl volumes were used for PCR, each with 2µl of HotShot extraction, 12.5µl MiFi taq (Bioline, UK), 8.5µl of molecular grade water, and 2µl of 10M primer (1µl of forward and reverse) (5′–TAAACTTCAGGGTGACCAAAAAATCA–3′). This included PCR blanks (negative controls) and single species positives (*Triops cancriformis*) (positive controls). PCR products were confirmed with gel electrophoresis on a 2% agarose gel stained with ethidium bromide, and commercially sequenced via the Sanger method (Sanger et al., 1977) using HCO2198 (Macrogen Europe, Amsterdam, Netherlands). Sequences were manually checked using CodonCode Aligner (v8.0.2) (Ma et al., 2015) (http://www.codoncode.com/). To remove PCR primers, the first 30bp of the reads were clipped off. Any poor-quality samples were discarded (shorter than 300bp or containing multiple ambiguous reads)(n=10). Nucleotide-nucleotide BLAST (BLASTN) (V1.13) (https://blast.ncbi.nlm.nih.gov/Blast.cgi?PROGRAM=blastn&PAGE_TYPE=BlastSearch&LINK_LOC=blasthome) was used to identify the species using existing sequences from Genbank to match with the cleaned SANGER sequences. The top hit of representative sequences was chosen (>95% match (identity), and query cover of >98% of input sequence) (Appendix 1). Instances where the manual identification did not match the top hit from BLASTN were considered manually misidentified.
